# Predatory selection of mucoid, antibiotic resistant *Pseudomonas putida* phenotype by myxobacterium *Cystobacter ferrugineus*

**DOI:** 10.1101/2020.11.06.371708

**Authors:** Shukria Akbar, D. Cole Stevens

**Author notes:** Address correspondence to D. Cole Stevens,.

## Abstract

Predation contributes to the structure and diversity of microbial communities. Predatory myxobacteria are ubiquitous to a variety of microbial habitats and capably consume a broad diversity of microbial prey. Predator-prey experiments utilizing myxobacteria have provided details into predatory mechanisms and features that facilitate consumption of prey. However, prey resistance to myxobacterial predation remains underexplored, and prey resistances have been observed exclusively from predator-prey experiments that included the model myxobacterium *Myxococcus xanthus.* Utilizing a predator-prey pairing that instead included the myxobacterium, *Cystobacter ferrugineus,* with *Pseudomonas putida* as prey, we infrequently observed surviving phenotypes capable of eluding predation. Comparative transcriptomics between *P. putida* unexposed to *C. ferrugineus* and the survivor phenotype suggested that increased expression of efflux pumps, genes associated with mucoid conversion, and various membrane features contribute to predator avoidance. The *P. putida* survivor phenotype was confirmed to be resistant to the antibiotics kanamycin, gentamicin, and tetracycline and to produce more alginate, pyoveridine, and phenazine-1-carboxylic acid than predator-unexposed *P. putida*. Unique features observed from the survivor phenotype including small colony variation, efflux-mediated antibiotic resistance, phenazine-1-carboxylic acid production, and increased mucoid conversion overlap with traits associated with *Pseudomonas aeruginosa* predator avoidance and pathogenicity. The survivor phenotype also benefited from increased predator resistance during subsequent predation assays. These results demonstrate the utility of myxobacterial predator-prey models and provide insight into prey resistances in response to predatory stress might contribute to the phenotypic diversity and structure of bacterial communities.

## Introduction

Abundant within soils and marine environments, predatory myxobacteria contribute to nutrient cycling within the microbial food web (1–4). Myxobacteria are prolific producers of antimicrobial specialized metabolites and display a cooperative, swarming predation strategy that can be readily reproduced and monitored within laboratory settings making them uniquely appropriate for assessment of predator-prey interactions (1, 3–5). Broadly considered generalist predators, myxobacteria capably predate a range of prey including both Gram-negative and Gram-positive bacteria as well as fungi (5, 6). Constitutive production of specialized metabolites and lytic proteins are often associated with this predatory range, and a variety of metabolites and enzymatic features have been reported to benefit predation (7–10).

However, relatively few examples of prey resistance to myxobacterial predation have been reported (11). Compared to features associated with bacterial strategies to avoid protozoan predators, prey avoidance of predatory myxobacteria remains underexplored (12–16). Examples of prey responses correlated with resistance to myxobacterial predation include *Escherichia coli* biofilm formation (17), *Bacillus subtilis* sporulation and production of bacillaene (18, 19), *Bacillus licheniformus* glycosylation of the predation-associated metabolite myxovirescin A (20), galactoglucan exopolysaccharide production and increased melanin production by *Sinorhizobium meliloti* (21, 22), and formaldehyde secretion by *Pseudomonas aeruginosa* (11). All of these features were discovered from predator-prey experiments utilizing the model myxobacterium *M. xanthus*.

We suspect the development of predator-prey pairings including myxobacteria other than *M. xanthus* might provide additional insight into prey resistance to myxobacterial predation. For our predator-prey experiments, the soil dwelling myxobacterium *Cystobacter ferrugineus* strain Cbfe23 was included due to a favorable growth profile and capability to quickly consume *P. putida* during standard predation assays (23–25). Also found within soils, root colonizing *P. putida* was chosen as prey due to an established ability to resist protozoan grazers (14). Initial predator-prey experiments provided a *P. putida* phenotype capable of avoiding *C. ferrugineus* predation. Herein we report the generation and predator avoidance of a *P. putida* phenotype resistant to myxobacterial predation using standard predator-prey experiments, differential gene expression data comparing the survivor phenotype with predator-unexposed *P. putida*, and traits observed to potentially contribute to predator avoidance.

## Results

### Selection of predation resistant *P. putida* phenotype

Utilizing predation assays where an inoculum of *C. ferrugineus* was introduced to the edge of an established spot of *P. putida* (24), we considered swarming overtaking the *P. putida* spot with no visible prey remaining to be an endpoint for assessment of remaining viable prey cells using colony forming unit (CFU) assays. Typically, *C. ferrugineus* swarmed predation sensitive *P. putida* with no visible prey after 3 days of co-cultivation on nutrient restrictive WAT agar (Figure 1A and 1D). However, from 100 total assays 5 provided minimally consumed and readily observable *P. putida* remaining after 10-14 days (Figure 1B and 1D). Additionally, instead of frontal attack at the predator-prey interface when exposed to predation sensitive *P. putida*, *C. ferrugineus* swarmed the perimeter of the survivor phenotype (Supplemental Figure 1). Exponentially fewer viable colonies of predation sensitive *P. putida* were observed from endpoint CFU assays (3-4 days for predation sensitive *P. putida;* 10-14 days for *P. putida* survivor phenotype) supporting the assumption that swarming resistance correlates with prey survival (Figure 1D and 1E). Also apparent from CFU assays, colonies of the survivor phenotype were consistently smaller when compared to predation sensitive *P. putida* (Figure 1C). Predation resistance of the survivor phenotype was also observed from additional predation assays with the optical density of prey inoculum doubled (OD_600_=1.0 versus OD_600_=0.5). (Supplemental Figure 2). Stocks of each *P. putida* survivor observed from predation assays were stored at −80°C as individual aliquots and utilized to determine features associated with the survivor phenotype that might contribute to predator avoidance.

**Figure 1:**
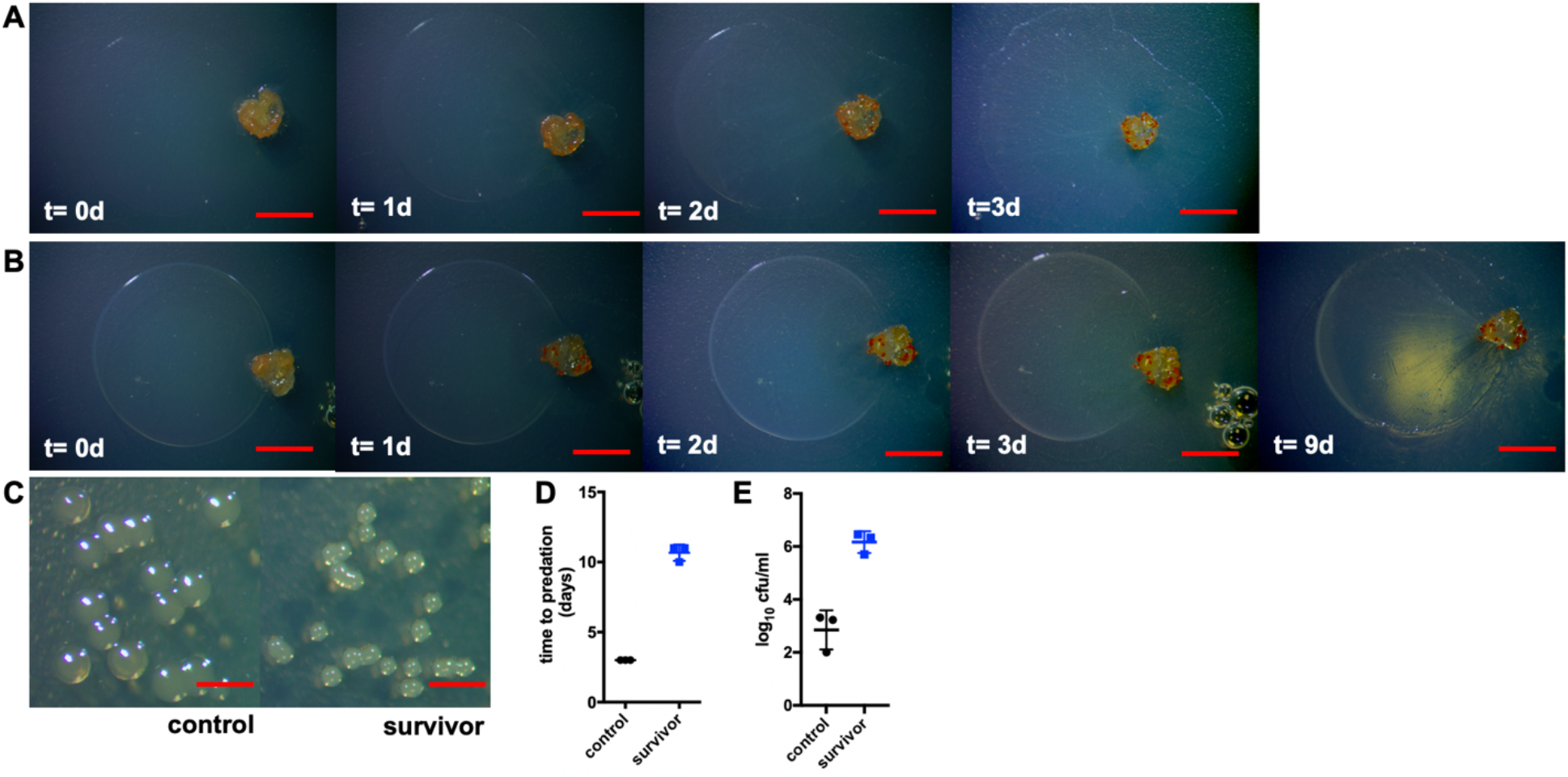
A) 4 day sequence of *C. ferrugineus* predation of *P. putida* with complete swarming of prey (t= d3). B) 4 day sequence depicting predator avoidance observed from *P. putida* survivor phenotype including an additional image from day 9 to demonstrate length of avoidance. C) Comparison of colony sizes of predator unexposed *P. putida* (left) and *P. putida* survivor phenotype (right) depicting small colony variance of the survivor phenotype. D) Time to predation data for *P. putida* control and *P. putida* survivor phenotype (n=3; p≤0.002) with E) end point colony forming unit data (n=3; p≤0.006) to determine differences in cell viabilities post-swarming. Unpaired t-test with Welch’s correction used for statistical analyses included in D and E; all scale bars depict 1mm.

### Differential gene expression of predator-unexposed *P. putida* and survivor phenotype

Utilizing RNA sequencing, we conducted comparative transcriptomic experiments between the predator unexposed (control) *P. putida* and the survivor phenotype (n=3 biological replicates with biological replicates for the survivor phenotype defined as an individually observed phenotype resistant to *C. ferrugineus* predation for ≥10 days). This comparative analysis determined that 1,178 genes were down-regulated ≥4-fold and 122 genes were up-regulated ≥4-fold across the survivor phenotype replicates when compared to control *P. putida* replicates (n=3; p≤0.05). Considering up-regulated genes from the survivor phenotype with the highest fold change values, apparent differences in overexpressed features specific to the survivor phenotype included genes involved in mucoid conversion, siderophore production, and a variety of membrane-associated proteins (Figure 2A). A 4 gene operon including a ferric uptake regulatory *(furA)* associated gene (WP_016501343.1), a fumarate hydratase (WP_016501342.1), a manganese superoxide dismutase *(sodA)* (WP_016501340.1), and a hypothetical protein with a conserved DUF2753 domain (WP_06501341.1) was significantly up-regulated in the survivor phenotype (Figure 2B and 2C). Expression of this operon from *P. aeruginosa* in response to iron limitation leads to elevated alginate production, an exopolysaccharide composed of mannuronic acid and guluronic acid monomers, and increased mucoidy (26, 27) and corroborates our observation of small colony variance from the survivor phenotype. Interestingly, the alginate regulatory elements MucA (WP_016498171.1) and MucB (WP_016498172.1) were both significantly down regulated in the survivor phenotype (Figure 2B). The anti-sigma factor MucA and the associated binding partner MucB inhibit mucoid conversion by sequestering the alternative sigma factor AlgU required for alginate production in *P. aeruginosa* (28–31). Increased transcription from the 4 gene, *furA* operon combined with decreased transcription of MucA and MucB, indicates that mucoid conversion might contribute to predator avoidance which parallels *P. aeruginosa* predatory avoidance mechanisms when exposed to grazing amoeba (32).

**Figure 2:**
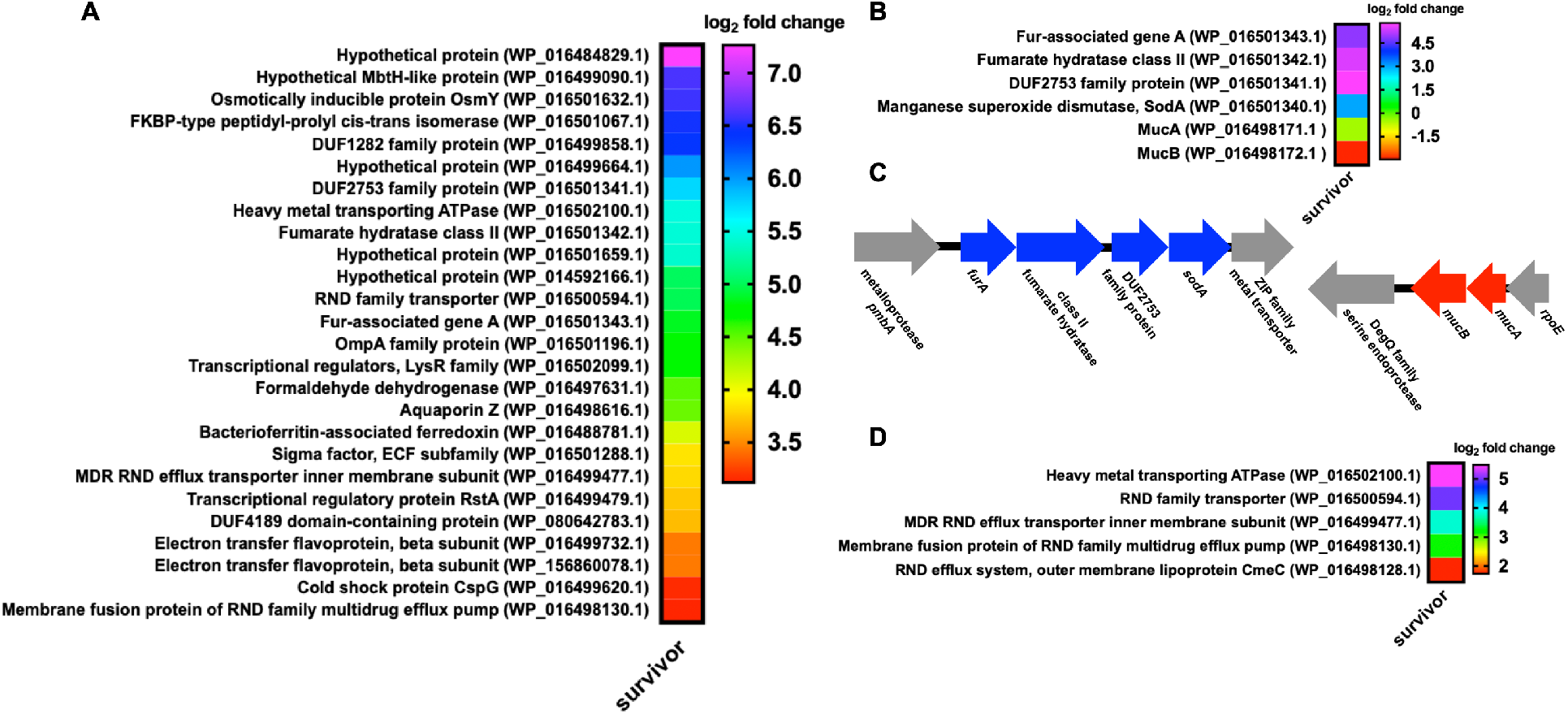
A) Most abundantly over-expressed genes from survivor phenotype when compared to predator unexposed *P. putida.* B) Up-regulated features associated with alginate production and the associated genomic context for each (C). D) Up-regulated transport proteins including those associated with antibiotic efflux. All data depicted as an averages from 3 biological replicates (p≤0.05).

Also associated with iron limitation, regulatory and biosynthetic genes contributing to production of the siderophore pyoverdine were significantly up-regulated in the survivor phenotype. These included PvdA an L-ornithine monooxygenase (WP_016499102.1) and PvdH a diaminobutyrate-2-oxoketoglutarate transaminase (WP_016498669.1) both involved in pyoverdine precursor biosynthesis, a regulatory transcription factor PvdS (WP_016498655.1), a proximal MbtH-like protein (WP_016499090.1), and a core nonribosomal peptide synthetase homologous to PvdL from *P. aeruginosa* (NP_746359.1) (Supplemental Figure 3) (33–36). Also associated with iron limiting conditions, a bacterioferritin-associated ferredoxin (WP_016488781.1) was upregulated in the survivor phenotype.

A variety of membrane-associated proteins were also overexpressed in the survivor phenotype including an osmotically inducible protein OsmY (WP_0016501632.1), an outer membrane protein A (OmpA) family protein (WP_016501196.1), an FKBP-type peptidyl-prolyl cis-trans isomerase (WP_0165010671), an aquaporin (WP_016498616.1), and a DUF1282 domain-containing hypothetical protein from the Yip1 superfamily (WP_016499858.1) (Figure 2A). While not a membrane associated feature, we also found the overexpression of a formaldehyde dehydrogenase (WP_016497631.1) by the survivor phenotype to be of note due to the recent observation that suggests formaldehyde secretion to be a predation-resistance trait of *P. aeruginosa* (11). The abundance of up-regulated genes that encode membrane-associated products in the survivor phenotype including those responsible for increased mucoid production provides insight into features involved in predation avoidance.

Numerous efflux and transport proteins were also among the most up-regulated genes observed from the survivor phenotype (Figure 2D). Upregulated efflux genes included inner (WP_016498130 and WP_016499477.1) and outer (WP_016498128.1) membrane components highly homologous to the AcrAD-TolC-type multidrug resistance-nodulation-division (RND) family of efflux pumps and the Mex RND efflux pumps known to facilitate *P. aeruginosa* resistance to aminoglycosides and tetracyclines (37–39). Genes encoding features homologous to multidrug efflux pumps contributing to virulence of human pathogens were also up-regulated in the survivor phenotype including a P-type ATPase (WP_016502100) and an additional RND/MmpL (Mycobacterial membrane protein Large) protein (Figure 2D). Afforded these observed differences in gene expression, we sought to determine if the survivor phenotype demonstrated antibiotic resistance, increased mucoidy, and increased production of the siderophore pyoverdine which have all been previously associated with predator avoidance strategies.

### Antibiotic resistance of survivor phenotype

Provided the observation that numerous transport proteins were overexpressed in the survivor phenotype (Figure 2D) including RND-type efflux pumps known to contribute to *P. aeruginosa* antibiotic resistances, we were interested to determine if the survivor phenotype exhibited antibiotic resistance. The survivor phenotype was capable of initially forming colonies on LB agar supplemented with the antibiotics gentamicin (10 μg/ml), kanamycin (50 μg/ml), and tetracycline (10 μg/ml), with no colonies from predation sensitive *P. putida* observed on identical medias. Subsequent growth curve assays comparing predation sensitive *P. putida* with the survivor phenotype when grown in LB and LB supplemented with antibiotics (Figure 3A and 3B) confirmed that the survivor phenotype was uniquely resistant to gentamicin, kanamycin, and tetracycline. These results combined with the overexpression of efflux-associated proteins observed from the survivor phenotype confirms that *C. ferrugineus* selection provides prey that benefit from antibiotic resistance and suggests efflux-mediated antibiotic resistance to be involved in *P. putida* response to predatory stress.

**Figure 3:**
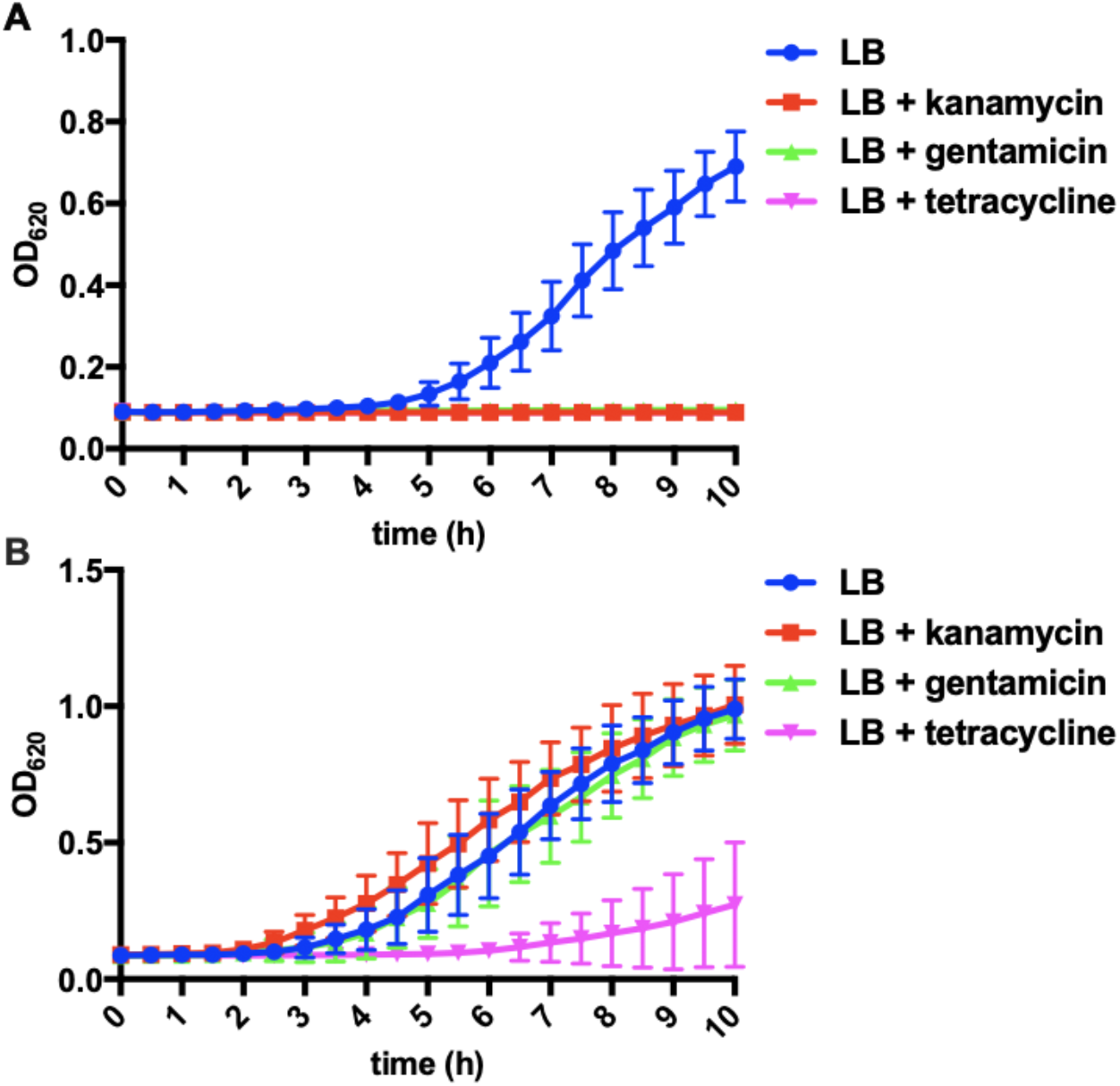
Growth curves for predator unexposed *P. putida* (A) and *P. putida* survivor phenotype (B) depicting antibiotic resistance unique to the survivor phenotype (n=12). LB supplemented with gentamicin (10 μg/ml), kanamycin (50 μg/ml), and tetracycline (10 μg/ml) where indicated.

### Increased production of alginate by survivor phenotype

Alginate-based formation of mucoid biofilms has previously been associated with *P. aeruginosa* avoidance of the protozoan grazer *Rhynchomonas nasuta* (32, 40–42). Utilizing an established carbazole assay as well as alginate antibody dot blots (43–45), we sought to determine if the increased transcription of genes known to impact alginate production observed from the survivor phenotype resulted in increased production of alginate when compared to predator unexposed *P. putida*. Comparing supernatants from predation sensitive *P. putida* with the survivor phenotype, we observed that the survivor phenotype indeed produced more alginate when compared to *P. putida* not exposed to *C. ferrugineus* (Figure 4).

**Figure 4:**
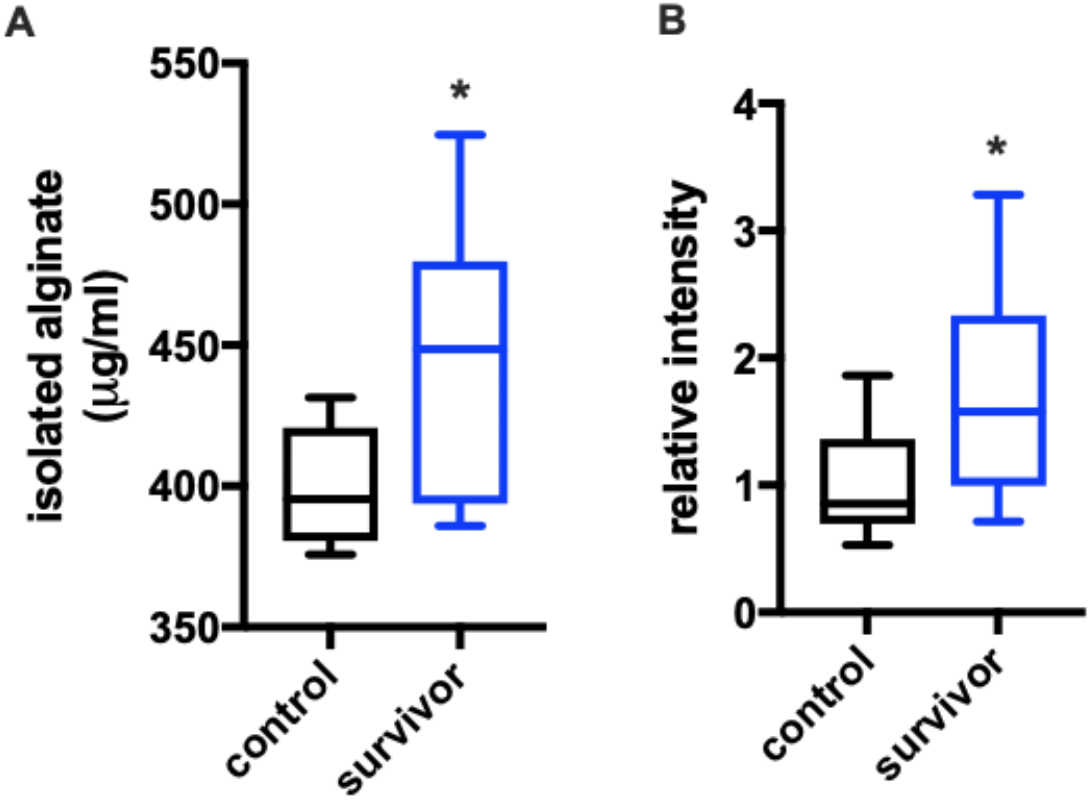
A) Isolated alginate concentration determined from carbazole assays depicting increased alginate production observed from the survivor phenotype (n=16; p≤0.02). B) Relative intensities of alginate dot blots generated from isolated alginate also depicting increased alginate production observed from the survivor phenotype (n=10; p=0.039). Relative units determined from average of control values with statistical significance calculated from an unpaired t test with Welch’s correction.

### Increased pyoverdine production by survivor phenotype

Significantly upregulated features associated with pyoverdine biosynthesis previously noted from the survivor phenotype in our transcriptomic data suggested that increased pyoverdine production might be an observable survivor trait. Fluorescence of extracellular pyoverdine was quantified from clarified medias of predator unexposed *P. putida* and the survivor phenotype using methodology established by Imperi *et al* (46, 47). Indicative of increased pyoverdine secretion, extracellular fractions from the survivor phenotype exhibited a 3-5 fold increase in fluorescence when compared to extracellular fractions from predator unexposed *P. putida* (Figure 5). Although the genome of the *P. putida* type strain includes a pyoverdine biosynthetic gene cluster, no structurally elucidated pyoverdines have been characterized from the *P. putida* type strain, and these results merely suggest the presence of fluorescent pyoverdine-like metabolites potentially produced by the strain. Regardless, these results corroborate overexpression of pyoverdine biosynthetic pathway components observed in our transcriptomic analysis and suggest that pyoverdine production may contribute to the predator avoidance trait of the survivor phenotype.

**Figure 5:**
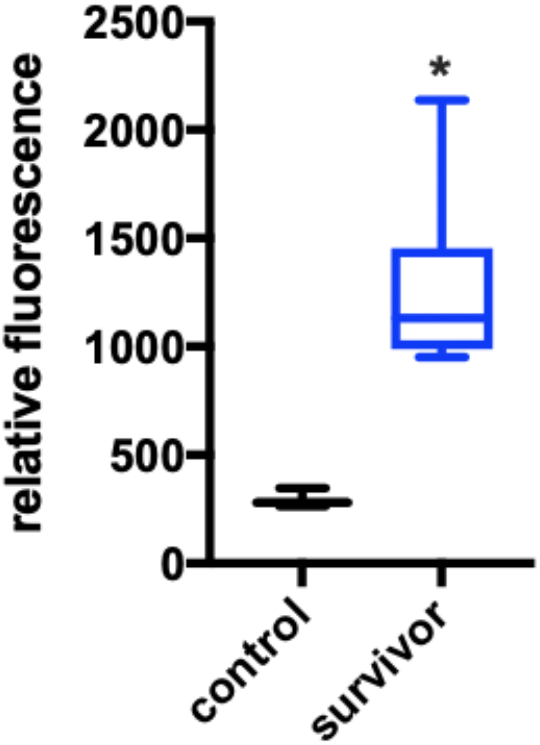
Relative fluorescence of extracellular pyoverdine depicting increased pyoverdine production from the survivor phenotype (n=12; p≤0.0001). Relative units determined from average of control values with statistical significance calculated from an unpaired t test with Welch’s correction.

### Increased production of phenazine-1-carboxylic acid by survivor phenotype

Untargeted mass spectrometry was used to assess additional differences in detectable quantities of *P. putida* metabolites comparing organic phase extracts from predator unexposed *P. putida* and the survivor phenotype. Initial analysis of resulting data using Global Natural Product Social Molecular Networking (GNPS) (48) resulted in a library hit for phenazine-1-carboxylic acid ([M+H]=225.066) that was only observed in extracts from the survivor phenotype (Figure 6A). Subsequent comparison utilizing XCMS-MRM (49) provided the statistical significance (n=3; p=0.0509) in detected phenazine-1-carboxylic acid between extracts from predator unexposed *P. putida* and the survivor phenotype (Figure 6B). Interestingly, phenazine production in *P. aeruginosa* contributes to redox maintenance during biofilm formation and promotes efflux-based antibiotic resistance (50–53). Despite no significant change in expression of putative phenazine biosynthesis genes *phzD* (SUD74861.1) and *phzF* (SUD71919.1) from our comparative transcriptomic data (54–57), we suspect that higher quantities of phenazine-1-carboxylic acid detected in extracts from the survivor phenotype may contribute to the increased transcription of efflux pump components and antibiotic resistance observed from the survivor phenotype. However, associations between phenazine production and efflux-mediated resistance have only been reported from *P. aeruginosa*, and any parallels potentially observed in our data require further investigation. Also of note, phenazine production has been observed to protect *Pseudomonas aureofaciens* from predatory myxobacteria (58). These results suggest that the survivor phenotype produces significantly greater quantities of phenazine-1-carboxylic acid when compared to predator-unexposed *P. putida*.

**Figure 6:**
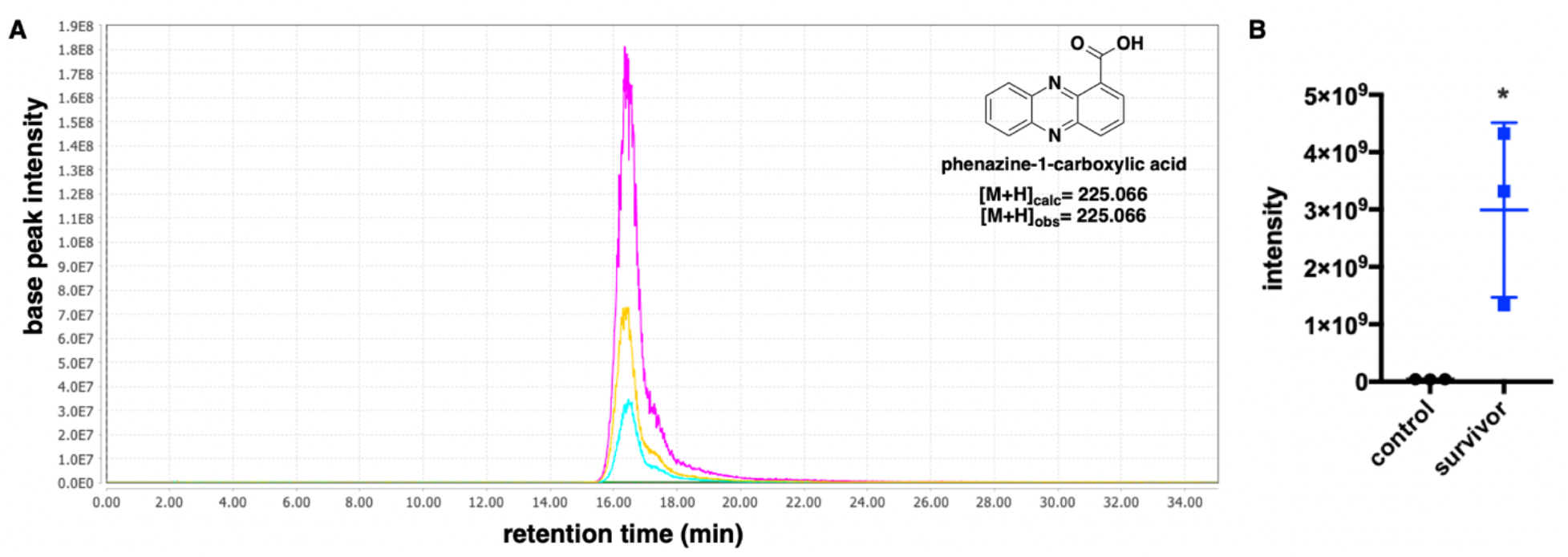
A) Extracted ion chromatograph (224.9-225.1 m/z) depicting presence of phenazine-1-carboxylic acid in extracts from survivor phenotype replicates (n=3; magenta, yellow, cyan) and absence in extracts from predator unexposed *P. putida* replicates (n=3; red, blue, green). Chromatograph rendered with MZmine v2.37. B) Comparison of detected phenazine-1-carboxylic acid in predator unexposed *P. putida* extracts (labelled control) and survivor phenotype extracts (blue). Box plot data and statistical analysis (n=3; p=0.0509) generated by XCMS v3.7.1 and figure rendered with Prism v7.0d.

### Predation resistance of survivor phenotype in subsequent predation assays

Additional subsequent predation assays were done to assess the predator avoidance trait of the *P. putida* survivor phenotype with *P. putida* not exposed to *C. ferrugineus* included as a control. These predation assays included an assay as previously described with predator cells directly introduced to the edge of established prey, a similar slightly modified assay with predator introduced at a distance of 5-7 mm from an established spot of *P. putida,* and a lawn culture assay (23) monitoring predator swarming diameters when placed at the center of prey embedded in agar. For direct contact assays, time to complete swarming was recorded. As observed in our prior predator-prey experiments, compared to control *P. putida* the survivor demonstrated significantly increased predator avoidance (Figure 7A). The survivor phenotype also demonstrated resistance to swarming compared to predation sensitive *P. putida* during identical direct-contact predation assays conducted on nutrient-rich VY/2 agar (Supplemental Figure 4) For indirect contact predation assays, time to predator-prey contact and time to complete swarming were recorded. Time to predator-prey contact was between 1-2 days for all samples, and as observed in the directed contact predation assays the survivor phenotype demonstrated increased predator avoidance when compared to control *P. putida* (Figure 7B and 7C). However, no differences between swarming diameters of *C. ferrugineus* grown on lawns of control *P. putida* and the *P. putida* survivor phenotype were observed (Figure 7D). These data suggest the *P. putida* survivor phenotype benefits from a predator avoidance trait when subjected to 2 out of 3 subsequent predation assays.

**Figure 7:**
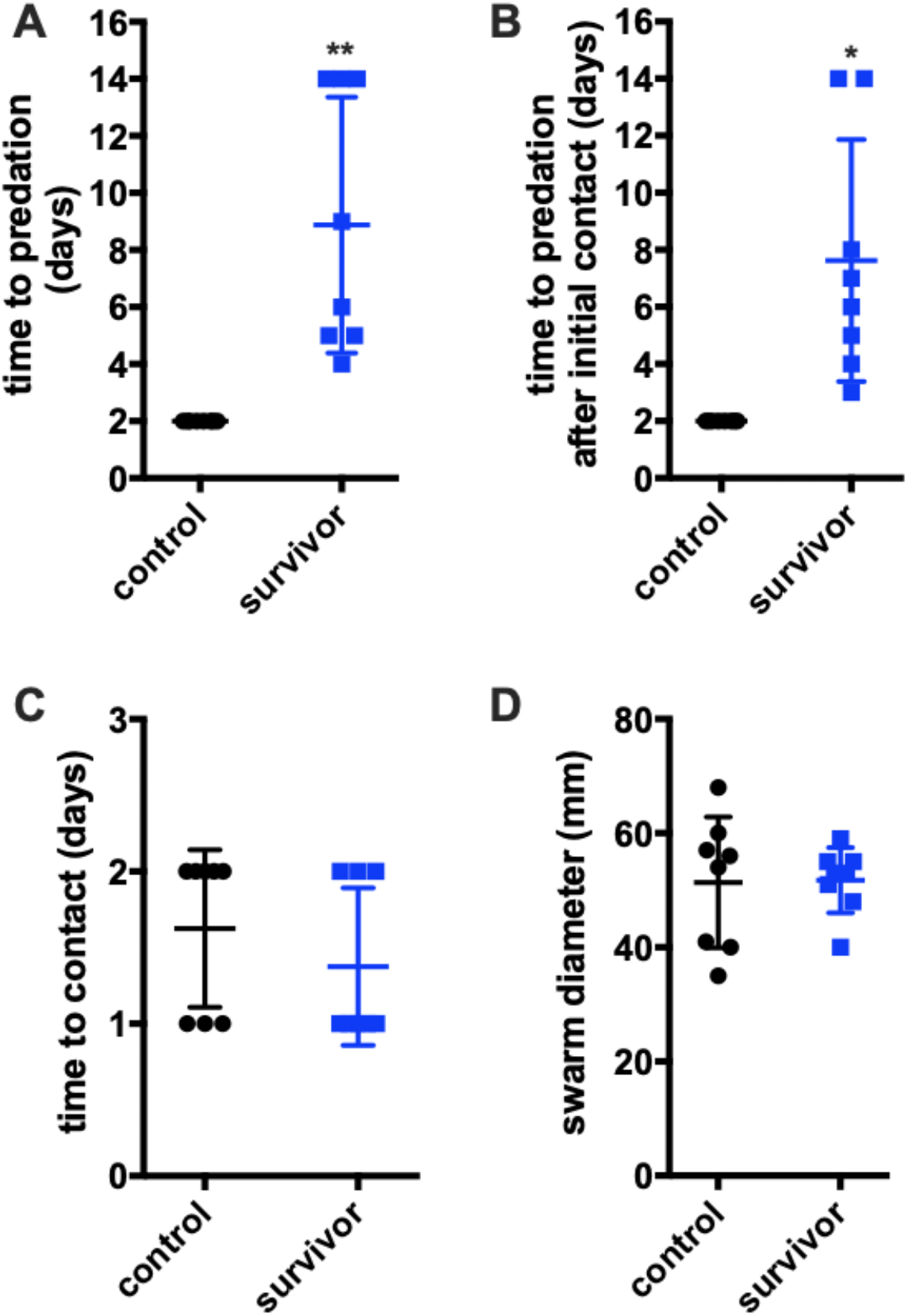
A) Direct contact predation assay as depicted in Figure 1 comparing control (predator unexposed) *P. putida* with freezer stocks of the survivor phenotype (biological replicates) (n=8; p≤0.0001). B) Modified indirect contact predation assay with prey introduced 2 cm from predator (n=8; p≤0.01) and time to predator-prey contact recorded (C). Swarming assays with *C. ferrugineus* plated on prey lawns and swarming diameters recorded after 7 days (n=8). Statistical significance calculated using an unpaired t test with Welch’s correction.

## Discussion

Building upon previously reported resistance strategies to avoid predatory myxobacteria discovered from predator-prey experiments (11, 17–22), predation experiments including *C. ferrugineus* and *P. putida* provided a prey phenotype significantly resistant to predation. Traits associated with prey resiliency during bacterial predator-prey interactions remain underexplored despite growing evidence that predatory stress contributes to the development of phenotypic traits associated with persistent clinical pathogens (12, 16, 59). As an example, clinical isolates of persistent *P. aeruginosa* from cystic fibrosis patients exhibit increased mucoidy and small colony variation as well as efflux-mediated antibiotic resistance (60–62). Other human pathogens that have demonstrated overlapping factors involved in both virulence and anti-predation strategies include *Legionella pneumophila, Escherichia coli, Vibrio cholerae, Campylobacter jejuni, Listeria monocytogenes, Mycobacterium leprae,* and *Yersinia enterocolitica* (12, 16). Overlapping factors thus far observed to be associated with both virulence and predator avoidance include biofilm formation (15, 32, 42), quorum signal (63) and siderophore (64) small molecule biosynthesis, toxin production (65–67), expression of transport proteins such as type III and type VI secretion systems (68, 69) and antibiotic resistance-associated efflux pumps (70, 71). While the impact of predatory selection on virulence factors have primarily been reported from predator-prey experiments with grazing amoeba, efforts to determine the influence of generalist predators on microbial communities have suggested that bacterial predators also drive adaptation of prey traits that overlap with virulence in bacterial pathogens (5). Recently, Nair *et al.* reported that during predator-prey interactions with *E. coli* prey, the predatory myxobacterium *Myxococcus xanthus* selects for 2 traits associated with virulence, mucoidy and increased expression of the outer membrane protease OmpT (5). Compared to features associated with bacterial strategies to avoid protozoan predators, prey avoidance of myxobacterial predators remains underexplored. Although predator-prey experiments that utilize myxobacteria with various prey such as *Bacillus subtilis* (9, 18, 19), *E. coli* (5, 7), *Sinorhizobium melilotti* (21, 22), *P. aeruginosa* (11), and *Streptomyces coelicolor* (72) have been reported, all myxobacterial resistances have been discovered from predatorprey experiments that included the model myxobacterium *M. xanthus*. Utilizing an alternative myxobacterium *C. ferrugineus* provides the opportunity to further compare predatory selection and adaptive prey avoidance strategies.

Comparative transcriptomics provided key features up-regulated in the *P. putida* survivor phenotype that overlap with traits associated with *P. aeruginosa* predator avoidance and pathogenicity. Of these, 3 putative RND family transporter components with high homology to the MexAB-OprM and MexCD-OprM efflux systems known to mediate *P. aeruginosa* antibiotic resistance were up-regulated in the survivor phenotype (38, 39). Growth curve analysis determined that the *P. putida* survivor phenotype was indeed resistant to the aminoglycosides kanamycin and gentamicin as well as tetracycline; susceptibility to all 3 of these antibiotics was observed from predator unexposed *P. putida*. Also of note, the multidrug efflux pump CmeABC serves as a survival mechanism for *C. jejuni* when engulfed by the predatory amoeba *Acanthamoeba polyphaga* (71).

Several features associated with mucoid conversion were also up-regulated in the survivor phenotype including a 4 gene operon responsible for elevated alginate production in *P. aeruginosa* and the regulatory transcription factors MucA and MucB (26–28, 30). Adaptive mucoid phenotypes have been previously observed in response to myxobacterial predation (5), amoebal grazing (40), macrophages (73), and lytic phages (74). Increased alginate production by the survivor phenotype was confirmed using established carbazole assays and anti-alginate dot blots. Alginate-based mucoid conversion and biofilm formation have previously been associated with *P. aeruginosa* avoidance of grazing amoeba and could also contribute to the small colony variation observed from our survivor phenotype (32, 40, 41). These results and the observed alternative predatory pattern of *C. ferrugineus* during predator-prey interactions with the survivor phenotype parallel previous observation of mucoid *S. meliloti* phenotypes avoiding *M. xanthus* predation (22). Additionally, upregulation of an OmpA family protein observed in the survivor phenotype might also correlate with mucoid conversion as *P. aeruginosa* phenotypes that upregulate expression of OmpA also overproduce alginate (75). These results provide clear overlap between features contributing to avoidance of myxobacterial and amoebal predation and also afford additional evidence that the coincidental selection of prey virulence factors reported from amoebal predatory stress (12) might also be observed from predator-prey interactions involving myxobacteria. Both efflux-mediated antibiotic resistance and mucoid conversion have also been attributed to virulence of clinical isolates of *P. aeruginosa* (29, 59, 60, 76).

Additionally, our data provided key areas for follow-up investigations to assess the chemical ecology of the predator-prey interface and its contribution to predator avoidance such as the increased production of pyoverdines and phenazine-1-carboxylic acid detected from the survivor phenotype. For example, up-regulation of the pyoverdine biosynthetic pathway and a bacterioferritin-associated ferredoxin as well as increased pyoverdine in extracellular extracts observed from the survivor phenotype suggests an iron-depleted predator-prey environment similar to the previously reported interaction between *M. xanthus* and *Streptomyces coelicolor* (72). Enhanced production of the primary myxobacterial siderophore myxochelin by *M. xanthus* contributed to an iron-restrictive environment and triggered production of the pigment actinorhodin from *S. coelicolor.* Further investigation focused on the chemical ecology of predator-prey interfaces might also shed light on the various membrane features observed to be up-regulated in the survivor phenotype.

The survivor phenotype was also assessed with subsequent predation assays to determine the resiliency of the predator avoidance trait. While comparative transcriptomic data provided insight into key up-regulated features potentially involved in prey survival, down-regulation of 1,178 genes in the survivor phenotype when compared to predator unexposed *P. putida* suggests potentially diminished survivor fitness. Despite this, the survivor phenotype maintained the predator avoidance trait in 2 out of 3 subsequent predation assays. While it is unclear why no significant predator avoidance was observed from lawn culture predation assays, we suspect the introduction of predator to the center of prey lawns as opposed to consistent introduction to the peripheral interface of prey potentially reduce any predator avoidance mechanisms. Predator avoidance observed from direct and indirect contact predation assays and the heterogeneity associated with differences between the growth and metabolism of peripheral cells and central cells of bacterial colonies supports this suspicion (77, 78). Perhaps belaboring, we also suggest that these results again highlight the importance of chemical exchange at the predatorprey interface. We anticipate that *P. putida* phenotypes resistant to myxobacterial predation continually subjected to such predatory selection will afford adapted phenotypes for further scrutiny of any resulting mutations and genotypes. Ultimately, these results demonstrate the utility of predator-prey experiments using lesser studied myxobacteria and provide insight into how predatory stress might contribute to adaptive phenotypic diversity of bacterial communities.

## Materials and Methods

### Bacterial Strains and Cultivation

The myxobacterium *Cystobacter ferrugineus* strain Cbfe23, DSM 52764 was employed as the predator. *Pseudomonas putida* type strain, ATCC 12633, and the discussed *P. putida* survivor phenotype resulted from predation assays with this *P. putida* type strain were included as prey. *C. ferrugineus* was grown on VY/2 solid (1.4% w/v agar, 0.1% w/v CaCl_2_ * 2 H_2_O, 0.5% w/v Baker’s yeast, 500 μM vitamin B12) media for 5-7 days. Luria-Bertani (LB) solid (1.5% agar) and liquid media were utilized for the cultivation of *P. putida.* Predation experiments were performed on WAT agar (1.5% w/v agar, 0.1% w/v CaCl_2_, 20 mM HEPES) plates. All bacteria were grown at 30 °C.

### Predation assays

Prey spot assay was performed as described by Seccareccia et al. (24). Briefly, *P. putida* prey was grown in LB liquid media on a rotary shaker (150 rpm) at 30 °C for 16-18 h. Next, the bacterial cultures were sedimented by centrifugation at 4,000 g for 15 min, and the sedimented cells were washed and resuspended in TM buffer (50 mM Tris, pH 7.8, 10 mM MgSO4) to an OD_600_ of 0.5. Then the WAT agar plates were spotted with 150 μl of the freshly prepared bacterial suspensions, and the spots were dried in a laminar flow hood. Finally, *C. ferrugineus* was inoculated at the edge of the prey spot. The assay plates were incubated at 30 °C for 14 days or until the full visible swarming of *C. ferrugineus* on the prey spot. The assay was performed with 100 replicates. Any prey spot with visible biomass of *P. putida* after day 14 was considered as survivor phenotype of *P. putida,* and five observed *P. putida* survivor stocks were acquired that remained visible after day 14 of the assay.

For subsequent predation assays, *P. putida* predation sensitive (control) and the survivor phenotypes were employed as prey. The direct spot assay was performed as mentioned above. Statistical significance of recorded time to predation reported in was calculated using an unpaired t test with Welch’s correction in Prism 7.0d. The subsequent directed contact predation assay was performed on VY/2 (1.4% w/v agar, 0.1% w/v CaCl_2_ * 2 H_2_O, 0.5% w/v Baker’s yeast, 500 μM vitamin B12) media as well, considered as nutrient-rich media. For indirect contact spot assay, *C. ferrugineus* was inoculated at a distance of 5-7 mm from the established *P. putida* control and *P. putida* survivor prey spots. Next, time *C. ferrugineus* took to reach the prey spots and the time of its swarming over the spots was recorded. Indirect predation assays were performed with 8 replicates per phenotype. Statistical significance of recorded time to predation reported in days and recorded time to initial contact between *C. ferrugineus* and prey reported in days was calculated using an unpaired t test with Welch’s correction in Prism 7.0d.

Swarming predation assays were based on established lawn culture methods (23). In brief, *P. putida* prey phenotypes were grown in LB liquid media for 16-18 h at 30 °C with shaking at 150 rpm. The cultures were then subjected to centrifugation at 4,000 g for 15 min, and the sedimented cells were washed and resuspended in TM buffer (50 mM Tris, pH 7.8, 10 mM MgSO4) to an OD_600_ 0.5. Next, 1 ml of the cell suspension was spread onto a 14 cm diameter WAT agar plate and dried to form a uniform prey lawn. Finally, *C. ferrugineus* previously cultured on VY/2 agar media was spotted onto the center of the prey lawn. The assay plates were incubated at 30 °C, and the expansion of the *C. ferrugineus* swarm was measured on day 4. Lawn culture predation assays were performed with 8 replicates per phenotype with replicates from each survivor stock included. All 3 predation assays were performed simultaneously for all replicates. Statistical significance of *C. ferrugineus* swarm diameters reported in millimeters on day 4 was calculated using an unpaired t test with Welch’s correction in Prism 7.0d

### CFU assays

The colony forming units (CFU) were calculated as described by DePas *et al.* (17). Briefly, after the full swarming of *C. ferrugineus* over the prey spot, the agar slab under the prey spot was excised and suspended in 2 mL phosphate buffer saline (PBS). Next, a dilution series from this solution was inoculated on the LB plates, and the plates were incubated at 30 °C for 18 hr. Then, CFU/mL was calculated from the viable colonies that appeared on the LB plates. Finally, the CFU/mL of the survivor was compared with CFU/mL of the unexposed *P. putida,* considered as a control. Statistical significance of all CFU assays was calculated using an unpaired t test with Welch’s correction in Prism 7.0d.

### RNA sequencing

Triplicate samples of *P. putida* control and *P. putida* survivor phenotype grown from stocks in LB at 37°C for 18 hr and pelleted via centrifugation were stored in *RNAlater* solution for sequencing. The DNA from *P. putida* type strain cells (200 μl) was extracted using MagAttract HMW DNA Kit (Qiagen). The DNA was eluted in 100 uL AE buffer. The concentration of DNA was evaluated (Supplementary Table 1) using the Qubit^®^ dsDNA HS Assay Kit (Life Technologies). The library was prepared using Nextera DNA Flex library preparation kit (Illumina) following the manufacturer’s user guide. 50 ng DNA was used to prepare the library. The sample underwent the simultaneous fragmentation and addition of adapter sequences. These adapters are utilized during a limited-cycle (6 cycles) PCR in which unique index was added to the sample. Following the library preparation, the final concentration of the library (Supplementary Table 1) was measured using the Qubit^®^ dsDNA HS Assay Kit (Life Technologies), and the average library size (Supplementary Table 1) was determined using the Agilent 2100 Bioanalyzer (Agilent Technologies). The library was diluted (to 6.5 pM) and sequenced paired end for 500 cycles using the MiSeq system (Illumina).

Total RNA was isolated from triplicate samples of the *P. putida* control, the *P. putida* survivor phenotype from stocks, and the kanamycin resistant phenotype using the RNeasy PowerSoil Total RNA Kit (Qiagen) following the manufacturer’s instructions. 400 μl cell sample was used for extractions. The concentration of total RNA was determined (Supplementary Table 2) using the Qubit^®^ RNA Assay Kit (Life Technologies). For rRNA depletion, first, 1000 ng of total RNA was used to remove the DNA contamination using Baseline-ZERO™ DNase (Epicentre) following the manufacturer’s instructions followed by purification using the RNA Clean & Concentrator-5 columns (Zymo Research). DNA free RNA samples were used for rRNA removal by using RiboMinus™ rRNA Removal Kit (Bacteria; Thermo Fisher Scientific) and final purification was performed using the RNA Clean & Concentrator-5 columns (Zymo Research). rRNA depleted samples were used for library preparation using the KAPA mRNA HyperPrep Kits (Roche) by following the manufacturer’s instructions. Following the library preparation, the final concentrations of all libraries (Supplementary Table 2) were measured using the Qubit^®^ dsDNA HS Assay Kit (Life Technologies), and the average library size was determined using the Agilent 2100 Bioanalyzer (Agilent Technologies). The libraries were then pooled in equimolar ratios of 0.6 nM, and sequenced paired end for 300 cycles using the NovaSeq 6000 system (Illumina). Differential expression between the resulting transcriptomes was calculated from pair-wise analysis of trimmed mean of M-values (TMM) normalized read counts (79) including fold change, counts-per-million (CPM), and associated p-values using the R-package, edgeR (80). Differential expression data with p-values ≤ 0.05 were considered statistically significant. Genome and RNA sequencing was conducted by MR DNA (Molecular Research LP). All fold change, CPM, and associated p-values resulting from differential expression experiments as described included as supplementary data files TABLE 1 (*P. putida* survivor compared with *P. putida* control). All raw RNAseq data is publicly available at the NCBI Sequence Read Archive (PRJNA577468).

### Antibiotic Susceptibility Assays

The standard growth curve assay was employed to determine the sensitivity of the unexposed *P. putida* and the survivor phenotype towards kanamycin (50 mg/mL), gentamicin (10 mg/mL), and tetracycline (10 mg/mL) antibiotics. The Lauria Bertani (LB) broth without any supplemented antibiotics was used as a control. The test was performed in a 96-well microtiter plate. In a 96-well plate, each well having 200 μL of LB or LB supplemented with an antibiotic was inoculated with a colony of *P. putida.* Next, the absorbance at OD_620_ was recorded using a plate reader every 30 min for 10 hr to examine the growth dynamics of each phenotype. The test was performed in 12 replicates per condition for both phenotypes. Recorded OD_620_ values were plotted using Prism version 7.0d with error bars depicting standard deviation of replicate data.

### Carbazole Assays

The alginate was isolated according to Jones *et al.* (81). In brief, both *P. putida* phenotype unexposed to the predatory stress and the survivor phenotype were grown overnight in 2 ml LB media and adjusted to an OD_600_ of 1.00. Following centrifugation, 1 ml of supernatant was treated with 2% cetyl pyridinium chloride to precipitate the alginate. The precipitated alginate was further collected by centrifugation. It was resuspended in 1 M NaCl and then re-precipitated in cold isopropanol. Precipitated alginate was suspended in 150 μl 0.9% (w/v) saline solution. Following established protocols (43, 44), a 50 μL aliquot of the isolated alginate and a dilution series of standard alginic acid (Sigma) were mixed with 200 μL of a solution of 25 mM sodium tetraborate in sulfuric acid and added in a 96-well plate. Next, the plate was heated for 10 min at 100 °C in an oven. After cooling at room temperature for 15 min, 50 μl of a 0.125% carbazole solution in absolute ethanol was added. Then, the plate was re-heated at 100 °C for 10 min in an oven and cooled down at room temperature for 15 min. Finally, the plate was read in a CLARIOstar microplate reader (BMG Labtech Inc., Cary, NC, USA) at a wavelength of 550 nm. Quantities of alginate per sample (μg/ml) were calculated from the resulting standard curve of purchase alginic acid, and statistical significance was calculated using an unpaired t test with Welch’s correction in Prism 7.0d.

### Anti-alginate Dot Plots

Alginate dot blots were generated using methodology from Lorenz *et al.* with slight modifications (45). Alginate was isolated as previously described, suspended in 150 μl 0.9% (w/v) saline solution, and 2 μl aliquots were pipetted onto a nitrocellulose membrane. After air-drying for 30 min, Tris buffered saline-Tween 20 (TBST) with 5% bovine serum albumin (BSA) was used for membrane blocking for 1 h. Membrane was then washed 3x for 10 min with TBST and subsequently incubated with the alginate antibody (Sigma, monoclonal, anti-mouse) overnight at 4°C. The following day, the membrane was washed 3x for 10 min with TBST, incubated with IRDye 800CW Goat anti-mouse (Sigma) for 30 min, rinsed 2x with TBST, and imaged on a LI-COR Odyssey. Average relative intensities for imaged blots comparing isolated alginate from predator unexposed *P. putida* with alginate isolated from the survivor phenotype (n=10) are reported with statistical significance was calculated using an unpaired t test with Welch’s correction in Prism 7.0d.

### Assessment of Extracellular Pyoverdine

Extracellular pyoverdine fluorescence was detected from supernatants following methodologies Imperi *et al.* and Barrientos-Moreno *et al.* (46, 47). Clarified supernatants from overnight cultivation of 200 μl LB cultures of predator unexposed *P. putida* and the survivor phenotype adjusted to an OD_600_ of 0.07 were generated by centrifugation (10,000 rpm, 15m). Fluorescence was recorded at 455 nm upon excitation at 400 nm using a CLARIOstar microplate reader (BMG Labtech Inc., Cary, NC, USA). Sterile LB was used as a negative control to assess background fluorescence. Average relative fluorescence for supernatants from predator unexposed *P. putida* and survivor phenotype cultures (n=12) are reported with statistical significance was calculated using an unpaired t test with Welch’s correction in Prism 7.0d.

### Metabolite Extraction and Analysis

For metabolomic analysis, the survivor phenotype and predator unexposed *P. putida* were cultivated on VY/2 agar plates as described in the predation assay. VY/2 agar plates with TM buffer were used as a negative control. The plates were incubated at 30°C for 14 days. After the incubation period, agar was chopped and extracted with ethyl acetate (EtOAC). The EtOAc extracts were dried in vacuo to produce crude extracts for LC-MS/MS analysis. The crude extracts from each condition were generated in triplicate and analyzed as previously described (82). LC-MS/MS generated data was converted to .mzML files using MS-Convert, and the GNPS platform (48) was utilized as a dereplication tool to look for the library hits within publicly available natural products libraries. For the statistical analysis, using XCMSonline (49), multigroup analysis with HPLC orbitrap default settings was employed. MZmine 2.53 was used to generate extracted ion chromatograms (83–85).

## Author Contributions

SA performed the experiments. DCS supervised the project. SA and DCS conceived the project, designed the experiments, and wrote the manuscript.

## Acknowledgements

The authors appreciate funding and support from the National Institute of Allergy and Infectious Diseases (R15AI137996) and the National Institute of General Medical Sciences of the National Institutes of Health (P20GM130460). The authors appreciate Scot E. Dowd at MR DNA, Molecular Research LP for assistance throughout sequencing services provided, Dr. Sandeep Misra Manager of the Analytical and Biophysical Chemistry Core associated with the Glycoscience Center of Research Excellence (GlyCORE; P20GM130460) for LC-MS/MS analysis, and Dr. Nicole Ashpole for her assistance with anti-alginate dot blot generation and imaging.

