## Supplemental Material for "Predatory selection of mucoid, antibiotic resistant *Pseudomonas putida* phenotype by myxobacterium *Cystobacter ferrugineus*"

**Supplementary Material**

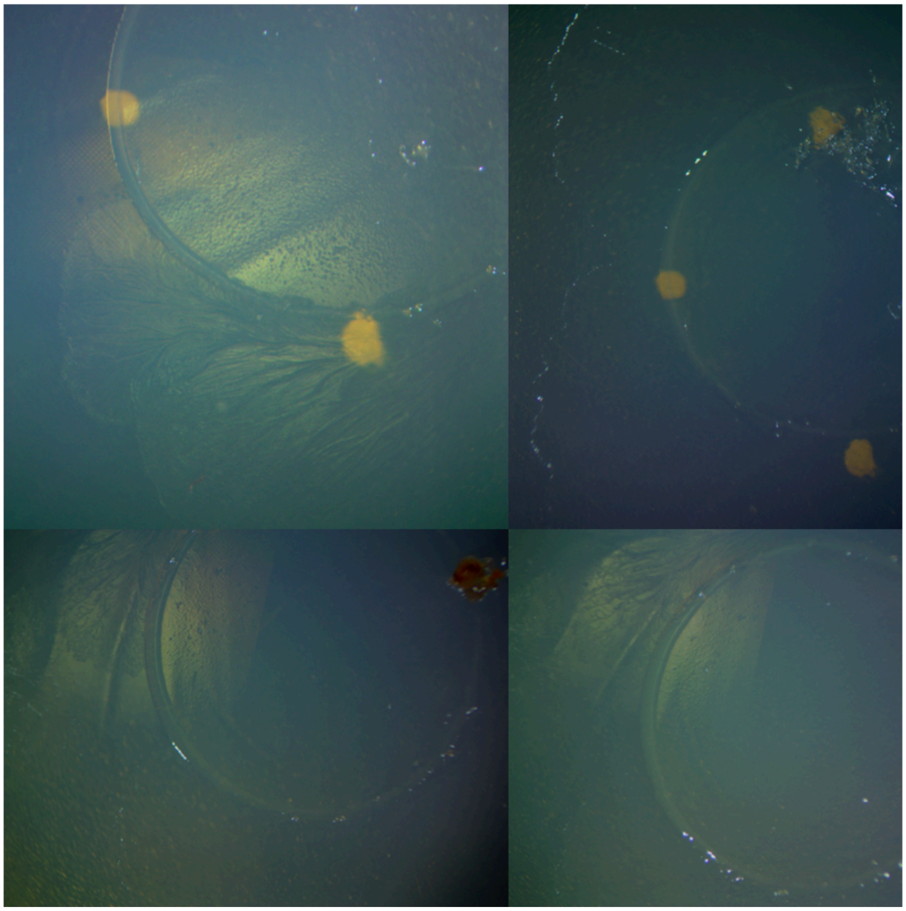

**Supplemental Figure 1:** Alternative swarming pattern observed from *Cystobacter ferrugineus* during predation assays with *Pseudomonas putida* survivor phenotype depicting perimeter swarming.

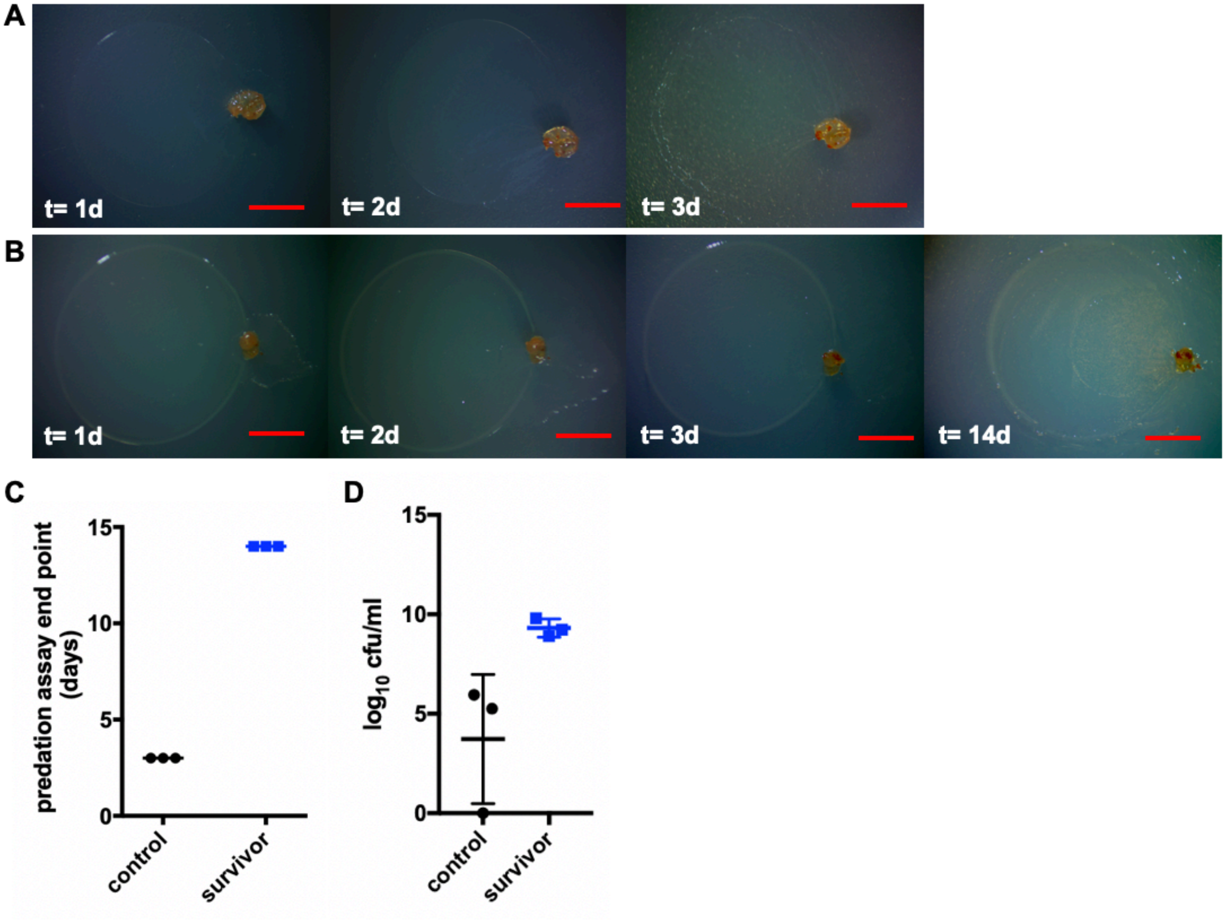


**Supplemental Figure 2:** Additional predator-prey assay with prey introduced at $OD_{600}$ of 1.0 A) 4 day sequence of *C. ferrugineus* predation of *P. putida* with complete swarming of prey (t= d3). B) 4 day sequence depicting predator avoidance observed from *P. putida* survivor phenotype including an additional image from day 14 to demonstrate length of avoidance. C) End point data for *P. putida* control and *P. putida* survivor phenotype predation assays (end point for survivor phenotype taken at 14 days for all samples as no complete swarming was observed) with E) end point colony forming unit data (n=3) to determine differences in cell viabilities post-swarming (no viable colonies observed from 1 control replicate). All scale bars depict 1mm.

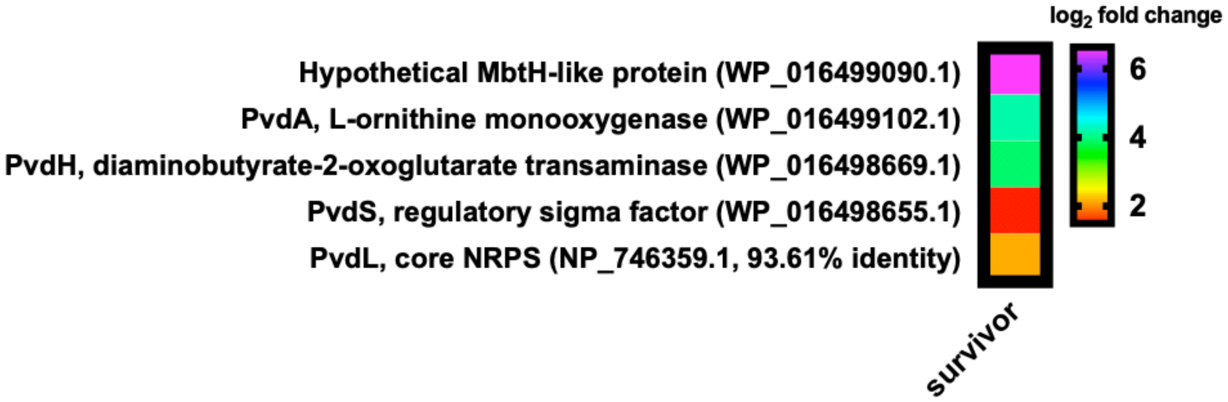


**Supplemental Figure 3:** Features associated with pyoverdine biosynthesis upregulated in the survivor phenotype. All data depicted as an averages from 3 biological replicates ($p \leq 0.05$).

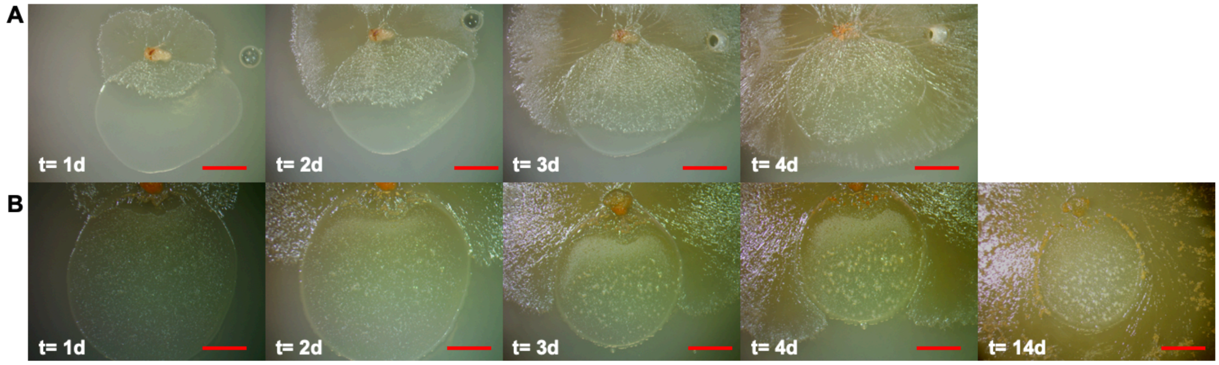


**Supplemental Figure 4:** Additional predator-prey assay on nutrient-rich VY/2 agar A) 4 day sequence of *C. ferrugineus* predation of *P. putida* with complete swarming of prey (t= d4). B) 4 day sequence depicting predator avoidance observed from *P. putida* survivor phenotype including an additional image from day 14 to demonstrate length of avoidance. Note the difference in swarming patterns with frontal swarming of control (A) and perimeter swarming of survivor phenotype (B).

| **Sample ID** | **DNA concentration (ng/µL)** | **Final library DNA concentration (ng/µL)** | **Average Library size (bp)** |
|---|---|---|---|
| PputidaTypestrain-3DNA | 34.0 | 18.20 | 901 |

**Supplementary Table 1:** Concentration of total DNA, final library concentration, and average library size for *P. putida* type strain sample used for genome sequencing.

| Sample | RNA Concentration (ng/uL) | Library Concentration (ng/uL) | Avg Library size (bp) |
|---|---|---|---|
| PputidaTypestrain-1 | 2360.0 | 39.60 | 610 |
| PputidaTypestrain-2 | 2360.0 | 37.60 | 591 |
| PputidaTypestrain-3 | 2360.0 | 43.80 | 634 |
| SurvivorPputida-1 | 2280.0 | 44.00 | 574 |
| SurvivorPputida-2 | 2560.0 | 32.60 | 435 |
| SurvivorPputida-3 | 1728.0 | 43.00 | 523 |

**Supplementary Table 2:** Concentration of total RNA, final library concentration, and average library size for *P. putida* type strain (predator unexposed control) and *P. putida* survivor phenotype.
